# Integrated tapered fibertrode for simultaneous control and readout of neural activity over small brain volumes with reduced light-induced artefacts

**DOI:** 10.1101/2020.07.31.226795

**Authors:** Barbara Spagnolo, Rui T. Peixoto, Leonardo Sileo, Marco Pisanello, Filippo Pisano, John A. Assad, Bernardo L. Sabatini, Massimo De Vittorio, Ferruccio Pisanello

## Abstract

Recognizing the neural patterns underlying different brain functions is essential to achieve a more comprehensive view on how small sets of neurons organize in complex 3D networks to determine different behaviours. In this framework, optogenetic techniques have been successfully proven as a powerful tool to control brain functions achieving millisecond temporal resolution and cell-type specificity, by combining the use of light-gated opsins and *ad-hoc* light delivery optoelectronic devices. However, targeting small brain volumes with simultaneous electrical recording results in the introduction of photoelectric artefacts, in particular when light emission and recoding sites are very close one to each other. In this work we take advantage of the photonic properties of tapered fibers to present a fully integrated fibertrode to target small brain volumes with abated photoelectric noise. The device hosts a light emitting window just below a recording pad, and exploits the angled light emission from the window to achieve simultaneous activation and electrical readout of small groups of cells with no photoelectric artifacts *in vivo*. Despite the highly non-planar surface of the fiber taper, window’s size, shape and electrode’s impedance can be modulated by controlling the fabrication parameters during focused ion beam milling and deposition, thus resulting in a versatile, integrated and customizable optogenetic tool for neurobiology studies in closed-loop configuration over small brain volumes.

## INTRODUCTION

The use of light to control the activity of genetically targeted neuron populations has introduced a new standard to study neural network functions. Its wide employment in neuroscience labs has prompted the development of novel optical devices capable of delivering light to the brain with configurations that vary in terms of ease of implantation, biocompatibility and light-delivery dynamics. As many of these optical probes integrate electrodes, they are capable of simultaneously controlling and recording neuronal activity and enable closed-loop control of specific neural populations with millisecond precision [1]–[5]. Real time feedback achieved by the combined use of optical stimulation and electrical readout of neural activity allows for a fully new paradigm in neuroscience, since the information provided can be used to go over the study of how different brain areas are connected and distributed. In fact, thanks to the bidirectionality of the information provided by closed-loop control of neural activity, it is possible to overcome causality studies and interfere with the ongoing brain events by normalizing an aberrant activation, inducing plasticity or consolidating memory [6].

One of the first optrode designs was based on flat-cleaved optical fibers glued on the top of implantable linear electrode arrays [7], [8]. Although prone to light-induced photoelectric artefacts due to direct illumination of the electrodes, this configuration still remains one of the most used, and recent studies have achieved low noise recordings by post-processing to correct for light-induced signals [9], [10]. However, the distance between the fiber and the recording sites results in an uneven and asymmetric distribution of light across the electrode array; moreover, this geometry illuminates large brain volumes compared to the size and pitch of the electrodes, often causing wide network activation beyond the local circuit of interest. Also, brain nuclei are often organized in small groups of cells where juxtaposed units play different roles, thus the use of large illumination volumes during optogenetic experiments can lead to unspecific activation of brain activity. To overcome these limitations and address small cellular groups for closed-loop optogenetic control, light sources should be integrated adjacent to recordings sites.

Several methods have been proposed to combine extracellular recording electrodes with local illumination. Solid state waveguides [5] and polymeric fiber optics [11] have been used to position light-delivery points in close proximity to electrode sites that have been processed with indium tin oxide [12], which removes high-frequency photoelectric artefacts and strongly mitigates low frequency light-induced noise. However, the stimulation volume remains large, comparable to that of standard fiber optics [13], [14]. Ridge waveguides that provide close stimulation and readout points with outcoupling gratings have also been employed [15], but require post-processing of signals to remove artefacts at light onset and offset. Implantable micro light emitting diodes (μLEDs) probes can also restrict illumination volumes, but the high current required for driving the emitter influences the recorded electrical signal, causing high frequency artifacts >50μV, limits the range of optical stimulus waveforms that can be applied [16]–[18], and the Lambertian emission profiles put constrains on the overall shape and size of the illuminated area [19], [20].

In this context, metal-coated microstructured tapered optical fibers (μTFs) have emerged as an effective system to deliver light to restricted brain volumes in superficial or deep brain areas. μTFs allow dynamic selection and modulation of light-emission properties [21], and the tapered tip reduces tissue damage. Light-emission can be restricted to small, micropatterned apertures fabricated along the fiber taper, emitting light along directions tilted with respect to the taper surface [22]. This peculiar feature of TFs is endowed by the photonic properties of the taper itself, which modifies the transverse wavevector of guided modes, generating light emission angles that would otherwise require the inclusion of light-redirecting elements [2], [23]–[25]. However, adding extracellular recording electrodes on the taper edge represents a significant technological challenge, because the small radius of curvature is incompatible with standard micro and nanofabrication planar techniques.

Here, we present a new design concept – the “fibertrode” – which integrates for the first time on a tapered optic fiber a platinum microelectrode very close (10 μm) to the light emitting site, yet exploits the peculiar photonic properties of the taper and the described fabrication strategies to eliminate photoelectric artefacts during optogenetic light trains. The taper acts upon the guided-mode wavevector to tilt the illumination pattern with respect to the long axis of the fibertrode, thus preventing direct illumination of the electrode as computationally confirmed by Monte Carlo simulations. The volume of stimulated tissue can be adjusted by changing the size of the FIB-milled optical window, while the precise Ion Beam Induced Deposition (IBID) process allows the fabrication of electrodes with adjustable sizes and impedances for different experimental needs.

We tested fibertrodes (in multiple window/electrode’s size configurations) in the mouse cortex and striatum of awake animals, showing spatially precise *in vivo* optogenetic activation and simultaneous artefact-free recording of local field potentials (LFPs) and extracellular action potentials. These results confirm that the angled emission assured by the light propagating properties of the tapered optic fiber, together with the FIB-IBID patterning of non-planar surfaces, allow obtaining optoelectronic devices to perform simultaneous optogenetic activation and electrical readout of neural activity over small brain areas avoiding the post-processing analysis usually implemented to remove photo-induced electrical artefacts.

## RESULTS

### Fibertrode design, fabrication and testing

A key challenge in realizing integrated fibertrodes is structuring the non-planar surface of the fiber taper, whose radius of curvature *r(x)* decreases along the waveguide axis *x* (see Figure 1A for definitions). This configuration requires (i) deposition methods that can achieve conformal insulation and metallization around a circular conical waveguide; and (ii) patterning techniques with spatial precision and resolution much higher than *r(x)*. The fibertrode was therefore designed exploiting the stack depicted in Figure 1A, with two aluminum (Al) layers alternated by two Parylene-C (Prl-C) insulating films, to obtain a metallic-confined waveguide [26], [27] and to prevents light leak onto the second Al layer. Al deposition was performed by continuously rotating the fiber during thermal evaporation with the fiber tip slightly tilted toward the crucible [28], resulting in a 600-800 nm thick coating. A conformal deposition of a 1μm-thick Prl-C film was then performed by physical vapor deposition, insulating the Al layer from the electrolyte [29] and creating a dielectric substrate for the realization of the recording pad. The second Al layer was then thermally evaporated without rotating the fiber, thus covering only one side of the waveguide and serving as electrical path for the microelectrode. This resulted in a conformal Al/Prl-C/Al stack, which was then further processed with focused ion beam (FIB) milling.

**Figure 1:**
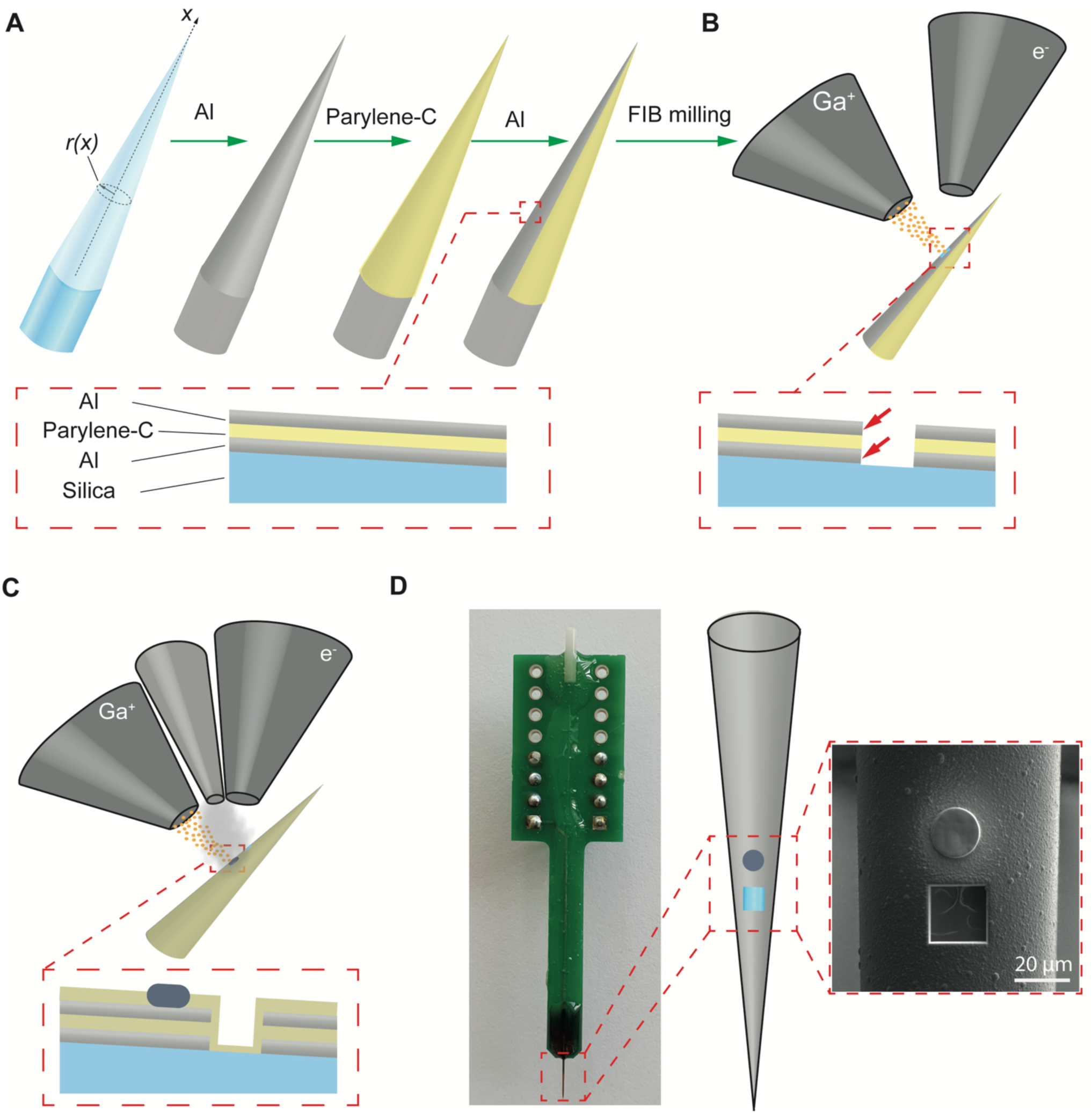
Fabrication process. (A) Two metal layers are deposited on a tapered fiber, interleaved by an insulating Parylene-C layer. (B) FIB milling is employed to open the optical window. (C) After depositing the final Parylence-C coating, FIB milling and FIBID are usedto realize the electrode. (D) Final device configuration.

Optical windows for outcoupling light were obtained by milling of the entire Al/Prl-C/Al stack over a defined area of customizable size and shape. The milling exposed the two Al layers along the window’s side walls (see red arrows in Figure1B), but a second Prl-C layer was then deposited to seal the device. To fabricate the electrode contact, the shallow Prl-C insulation was locally removed with FIB milling to expose the Al layer (Figure 1C). Then, Pt was slowly deposited via IBID in the recess, at a rate of ~0.45 μm^3^/sec. Platinum deposition was performed by scanning the ion beam on a circular surface slightly bigger than the recess on the second Prl-C layer, and the deposition parameters were optimized, accordingly to [28], to maximize texture homogeneity throughout the entire electrode pad. The final configuration of the device is displayed in Figure 1D.

Figure 2A displays typical IBID-deposited electrodes of diameters 7.5μm, 15 μm and 30 μm. Integrity and impedance of the fabricated electrodes were evaluated by Electrochemical Impedance Spectroscopy (see Materials and Methods). Independently from electrode size, a constant phase element (CPE) behavior was observed, with impedance phase close to 70° in the frequency range 1 Hz-100 kHz (Figure 2B).

**Figure 2:**
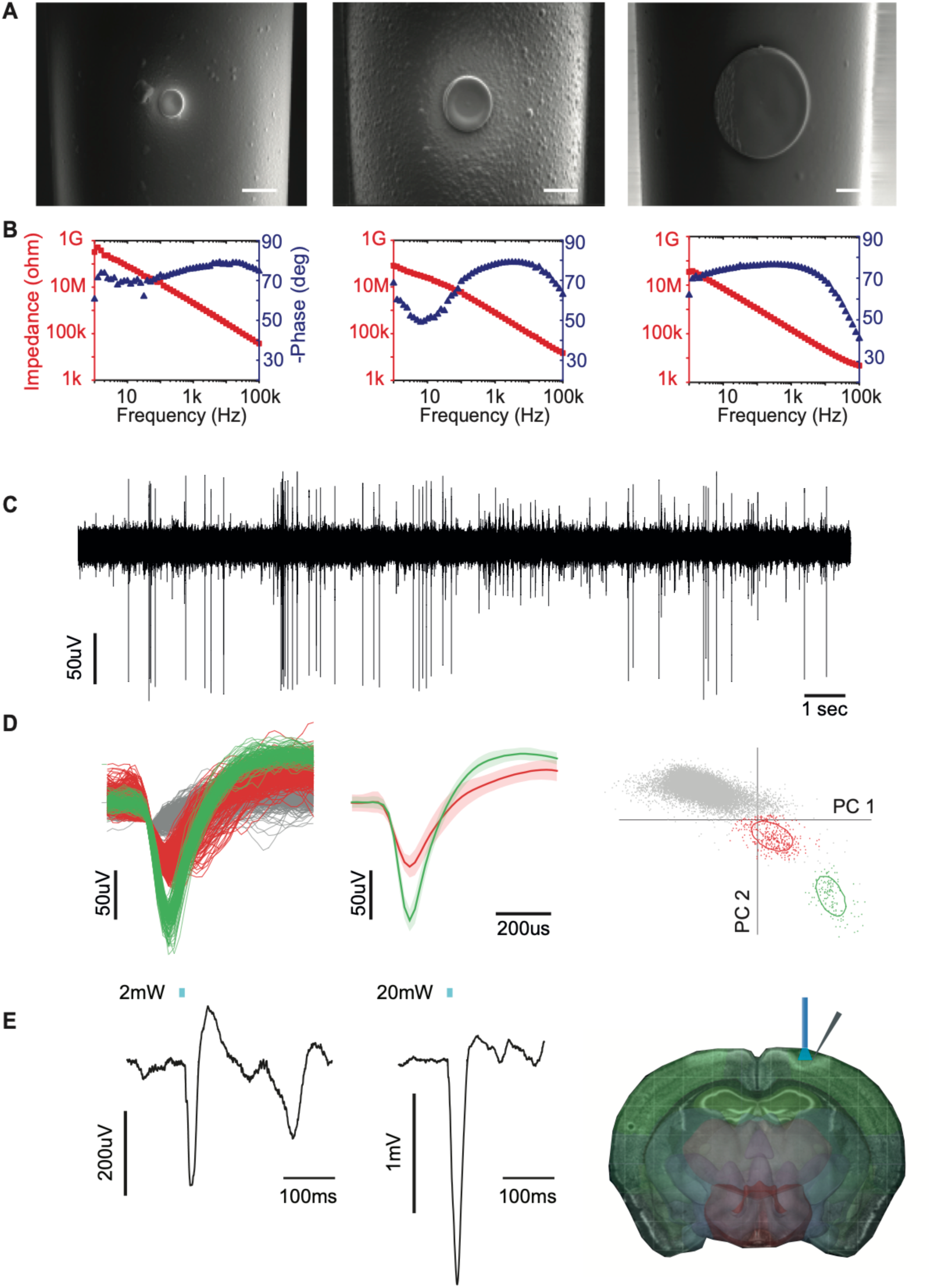
Electrical characterization and Extracellular recordings obtained with FIBID electrodes. (A) SEM micrographs of Pt electrodes with different diameters (7.5μm, 15μm and 30 μm). Scalebars represent 10μm. (B) Graphs showing impedance and phase val values measured at different frequencies for the three electrodes sizes. (C) Representative time trace of high-frequency recording in the frequency range 1KHz-10kHz with a 15μm electrode. D) Single-unit spike sorting, from left to right: sorted spikes, WT mouse n=1, unit1 (green)=138 spikes, unit2 (red)= 410 spikes; total waveforms 18387, Multivariate ANOVA 2D cluster space F(4,21088)= 16.3217, p=2.34 x10^-13 (average pm standard deviation of sorted spikes, principal component analysis. (E,left) Representative local field potentials (frequency range 300Hz-1kHz) recorded with the same electrode employed in Panels C and D. LFP recording of Thy1-ChR2 cortex. Traces represent average of n=5 and n=3 stimulation with 2mW and 20mW light, respectively (1 representative recording in 1 mouse). Averaging was performed in IgorPro using raw data (no post filtering or baseline subtraction). (E,right) Schematic representation of the experimental configuration. Light is delivered on top of the *dura mater* while recording neural activity in cortical layers 2/3.

As expected, impedance increased as the electrode sizes decreased, with typical values (measured at 1 kHz) of ~2.1MΩ, ~800kΩ, and ~150kΩ, for 7.5 μm, 15 μm, and 30 μm diameter electrodes, respectively. We then characterized their electrical performance *in vivo*, by recording spontaneous neural activity in layer 2/3 somatosensory cortex of non-anesthetized C57BL/6 mice. We employed tapered fibers covered with a single Al-Prl layer stack. Broadband recordings in the 300 Hz-10 kHz band revealed extensive neural spiking (Figure 2C), similar to recordings acquired with commercially available silicon-based microelectrode arrays. Offline waveform analysis of action potentials allowed sorting of multiple putative units (Figure 2D). These results validate the use of IBID-deposited electrodes for *in vivo* recordings of neuronal action potentials.

In addition, we recorded local field potentials (LFPs; 1-100Hz frequency band) in layer 5 (L5) of the somatosensory cortex of Thy1-ChR2 transgenic mice, using the configuration in Figure 2E(right): light was delivered extra-cranially with a 200 μm-core optic fiber placed over the *dura mater* near the site of the fibertrode implant. Optogenetic stimulation (488 nm laser light at 2mW or 20mW) was used to activate L5 pyramidal neurons dendrites that extend to superficial cortical layers. Stimulation elicited robust LFPs that were time-locked to the light pulses (Figure 2E, left).

Fibertrode’s light-emission properties in terms of emitted power density and light-output angles were characterized, and light-emission patterns from FIB-milled windows of varied dimensions are shown in Figure 3A. Light was launched into the fiber with a specific angle θ_in_ relative to the back facet of the fiber, to maximize output power density (Figure1C) and results show that at a taper diameter of 90 μm, the maximum output power was found at θ_in_~24°, for all windows sizes (Figure 3D) [22], [26].

**Figure 3:**
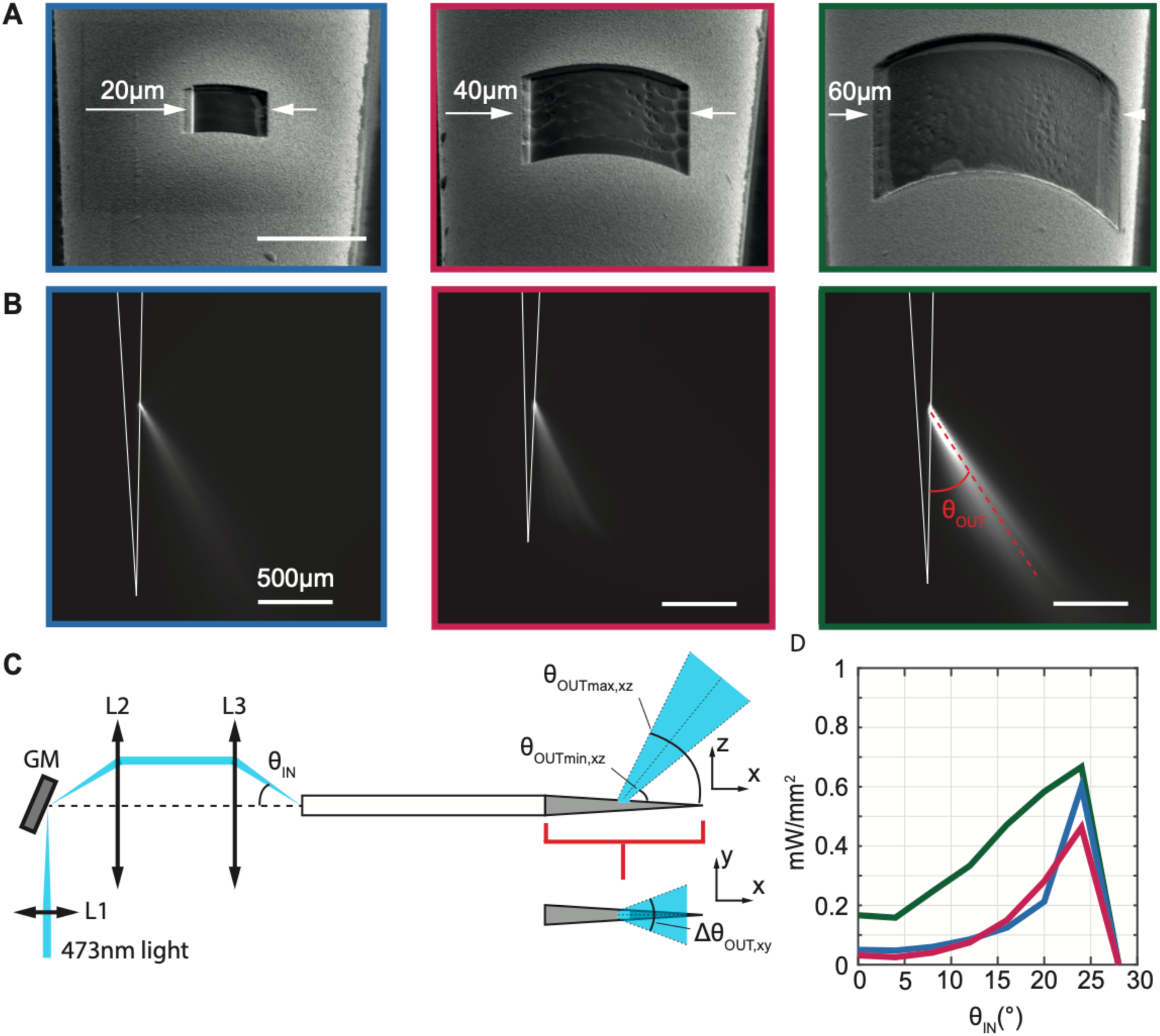
Light emission properties. (A) SEM micrographs of the realized optical windows (from left to right sizes are 20×20μm^2^, 40×40 μm^2^ and 60×60 μm^2^). Scalebar represents 30μm. (B) Emission patterns characterized in a 30mM fluorescein:PBS bath for each window size, at input angle maximizing emission intensity. Experimental output angle θ_out_ is ~28° the investigated window sizes. (C) Schematic of the light injection setup used for in vitro experiments. GM is a galvanometric mirror used to deflect the beam. Lens L1 focuses the beam on the GM, lens L2 collimate the beam while lens L3 focuses light onto the fiber core with a specific input angle. (D) Power density values measured at the window exit for different θ_IN._

In terms of outcoupling efficiency, a 2mW input into the taper resulted in an average output power density of 0.5-0.8mW/mm^2^. Importantly, the emitted light forms an angle with respect to the taper axis, resulting in a specific emission direction that we exploited to reduce direct illumination of the recording pad. This angle (θ_OUT;_ Figure 3C), has a value of ~28° independently from the window’s size. To estimate the amount of illumination received by the electrode, we implemented Monte-Carlo simulations to model the local scattering of light [30]. 1000 photons packets, each made of 1×10^5^ photons, were generated from a uniform distribution in the angular ranges θ_OUTmin,xz_ <θ_OUT,xz_ < θ_OUTmax,xz_ and -Δθ_OUT,xy_/2<θ_OUT,xy_< Δθ_OUT,xy_/2, in the xz and xy planes, respectively (see definitions in Figure 3C). Brain tissue was modeled with an Henyey-Greenstein scattering function, with parameters *n* = 1.360, *l* = 90,16μm, *g* = 0.89, *T* = 0.9874. Three-dimensional iso-intensity surface maps of the simulated emission patterns are shown in Figure 4A.

**Figure 4:**
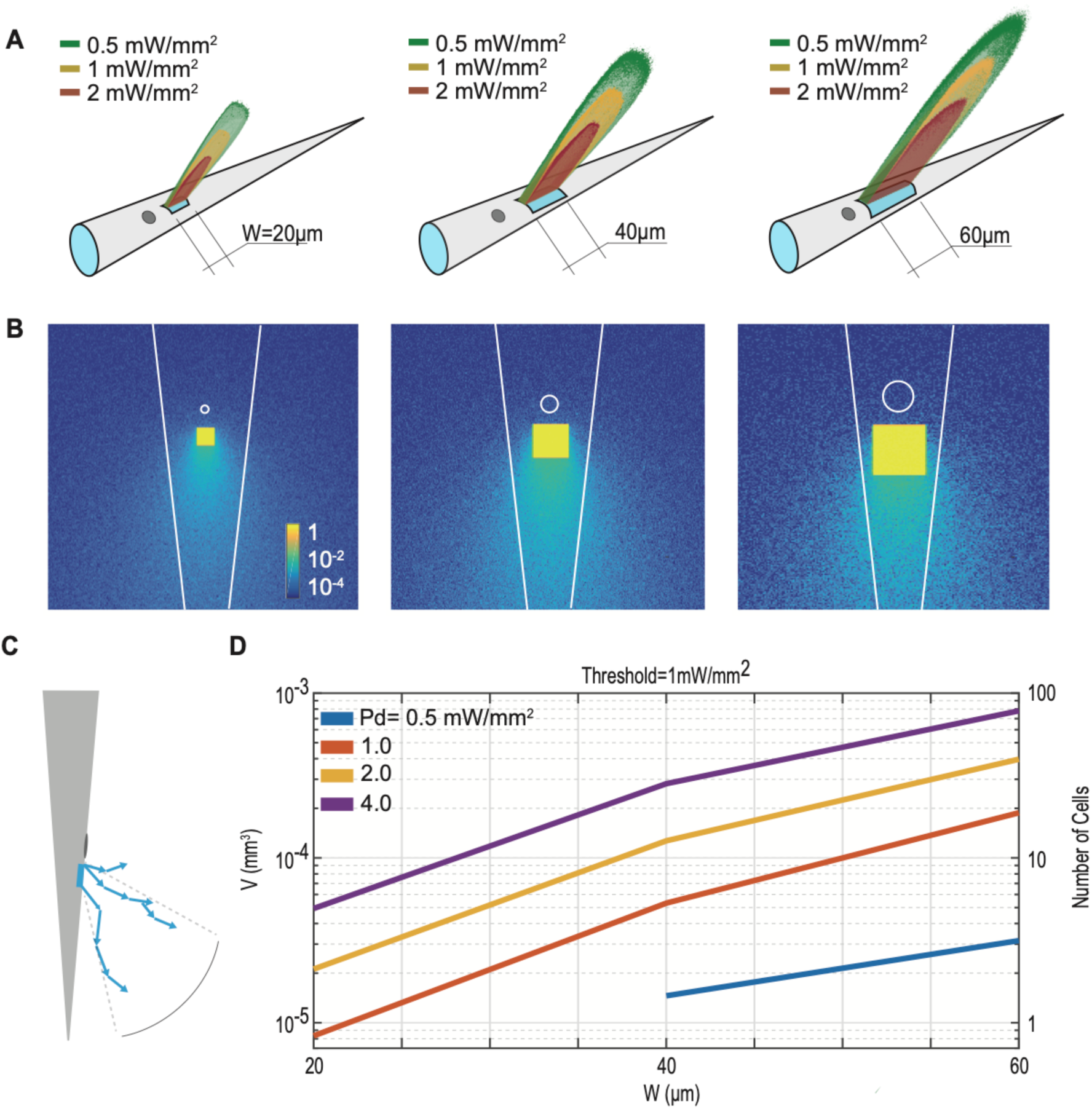
Monte-Carlo simulations. (A) Iso-intensity surface maps at 0.5 mW/mm^2^, 1.0 mW/mm^2^, and 2.0 mW/mm^2^ of the simulated light emission distribution for three different windows sizes, simulated by a Monte-Carlo approach. Data are shown for total output power from the windows of 2 mW Power density distributions at the fiber surface. Color scale is logarithmic. (C) Schematic representation of photons propagation at the window exit: electrode area is located in the region of lowest power density and is not interested by backscattered light. 1000 photons packets, each made of 10^5^ photons, were generated from a uniform distribution in the angular ranges θ_OUTmin,xz_ <θ_OUT,xz_ < θ_OUTmax,xz_ and -Δθ_OUT,xy_/2<θ_OUT,xy_< Δθ_OUT,xy_/2 in the xz and xy planes. For the three window’s size simulations were run with θ_OUTmin,xz_ = 25°, 21° and 27°, θ_OUTmax,xz=_ 42°, 38° and 33° and Δθ_OUT,xy_ of 18°, 24° and 30° for 20μm, 40μm and 60μm windows respectively. Brain tissue was modeled with an Henyey-Greenstein scattering function, with parameters *n* = 1.360, *l* = 90,16μm, *g* = 0.89, *T* = 0.9874. D) Estimation of volume enclosed by iso-surfaces at different power densities and different window sizes. Activation threshold is set at 1mW/mm^2^. Number of neurons related to the volumes are determined assuming a 10^5^ neurons per mm^3^.

Top-views of the fiber surface are shown in Figure 4B in logarithmic scale. The power-density distributions around the aperture were minimized above the optical window (Figure 4C), independent of the size of the aperture. The simulations suggest that window size can therefore be engineered to obtain the optimal stimulated volume for the specific application, without involving any direct illumination of the electrode. Indeed, considering the activation threshold for Channelrhodopsin-2 is 1mW/mm^2^ [31], from our simulations the stimulated tissue volume can be tailored from ~10^−5^ to ~10^−3^ mm^3^ by changing emitted power from 0.5 mW/mm^2^ to 4mW/mm^2^ for square windows of 20μm, 40μm and 60μm (Figure 4D). Assuming a density of 10^5^ neurons/mm^3^ in mouse cortex [32], these volumes correspond to ~1 to 100 neurons.

However, the two Al layers of the fibertrode could also be a source of photoelectric artefacts, due to light interaction with the inner layer. In a metallic-confined waveguide, both propagating and evanescent modes generate a surface current density on the metallic cladding [33], potentially giving rise to spurious electrical signals. In addition, electric currents can be generated by multi-photon absorption of 473nm light [9]. To characterize the presence of photo-induced electrical artefacts, we submerged the fabricated fibertrode in Phosphate Buffered saline Solution (PBS) and applied trains of long light pulses at 473 nm (200ms duty cycle 50%, power density 2mW/mm^2^, and input angle 24°), while recording voltage changes (Figure 5A). Long light pulses are not employed in typical optogenetic experiments, but were chosen as a worst-case test of device performances. The fibertrode design that we tested (double Al/Prl-C stack, 15μm electrode and 20μm optical window) showed a small light-dependent voltage change in the low-frequency band (LF: 3-250 Hz) (Figure 5A, upper panel) and no artefacts in the high-frequency band (HF: 0.25-10 kHz) (Figure 5A, lower panel).

**Figure 5:**
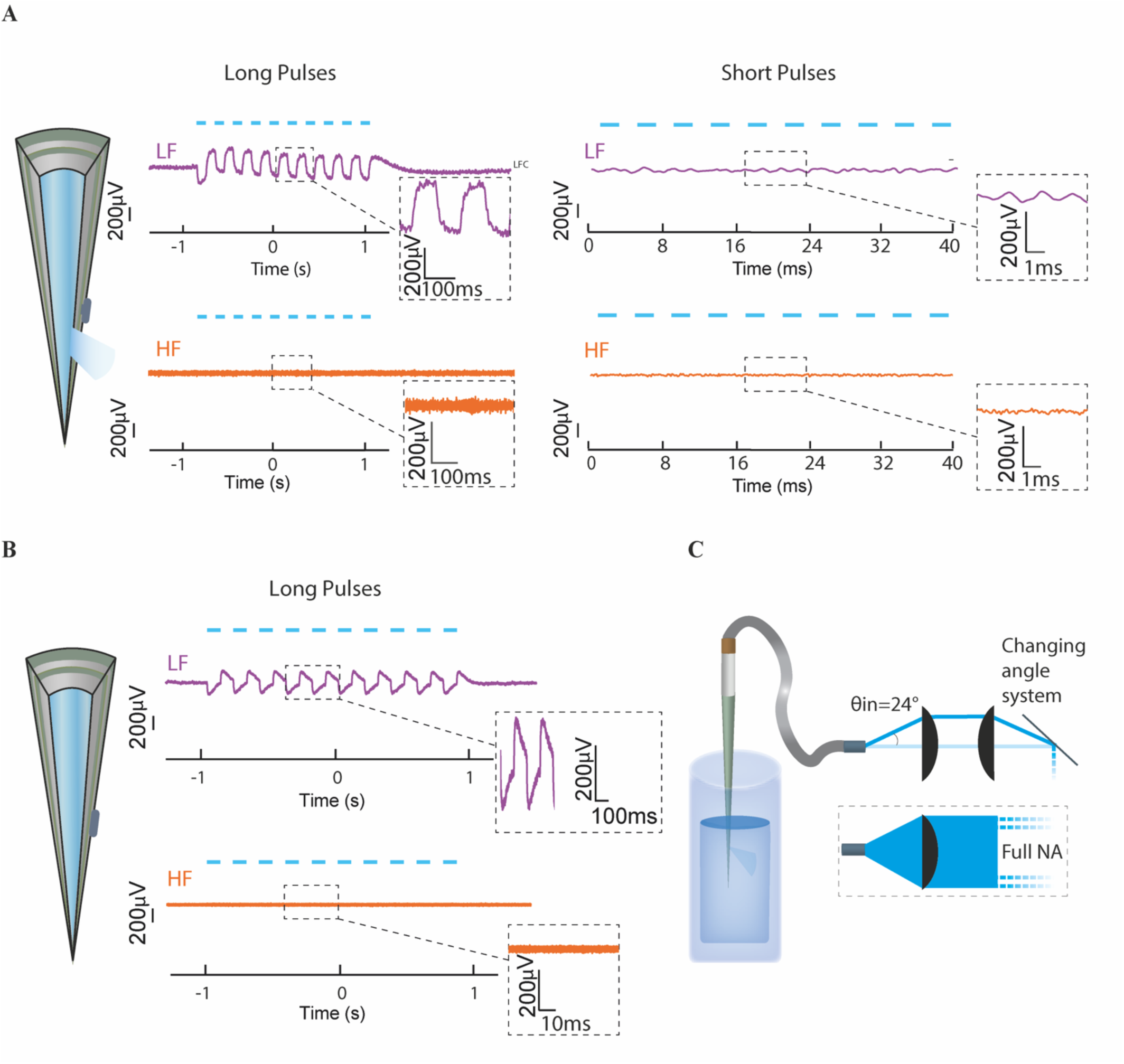
Light-induced photoelectric noise in PBS for low frequency (LF) and high-frequency (HF) channels. (A) Optimized fiber optrode tested with both 200ms and 2ms light pulses, obtained injecting light at an input angle θ_IN_=24°. (B) Optimized fiber optrode without optical window tested with 200ms pulses and full-NA input light, in a “worst-case” condition. (C) Schematic representation of the angled and full NA light injection paths used to fit the conditions of panel A and B respectively.

To test whether the optical window was responsible for the small LF optical artefacts, we also tested an analogous probe with no optical window (Figure 5B). As a worst-case test, we injected 200msec light pulses (duty cycle 50%) into the entire acceptance angle of the windowless fibertrode, to maximize temporal and spatial light interaction with the metal. Optical artefacts with the windowless probe (Figure 5B) were no worse than the windowed probe (Figure 5A), while the first Al layer with thickness >500nm should effectively shields the shallower Al layer hosting the electrode. A schematic representation of the angled and full NA optical paths is shown in Figure 5C.

The low frequency (LF) artefact could therefore arise from light interaction with the internal layer, generating a displacement current *I*_*D*_ in the external Al layer through capacitive coupling, which is then recorded by the amplifier. If so, *I*_*D*_ only generates an effect in the LF band; no artefacts were detectable in the HF spike band.

Importantly, when the laser pulse-width was reduced to a duration of 2ms, LF artefacts were not detectable (Figure 5A right). Taken together, these results show that optimization of the multi-layer fabrication process, together with IBID electrode fabrication and reciprocal positioning of the optical and electrical elements, leads to a marked reduction of photo-electrical artefacts in saline solution.

### In vivo optical control and readout of neural activity on small volumes without photoelectric artefacts

To determine whether the combined electrical and optical properties of fibertrodes are suitable for optogenetic stimulation and simultaneous monitoring of neural activity, we performed recordings in the striatum of awake Adora2a-Cre;Ai32^f/f^ mice expressing Channelrhodopsin2-EYFP (ChR2) in striatal spiny projection neurons of the indirect pathway (iSPNs).

Optical stimulation using a 473 nm laser (output power of 0.4-10mW/mm^2^) produced pronounced increases of spiking activity in neighboring neurons. Effective neural stimulation was obtained with all window sizes tested (20, 40 and 60 μm, Figure 6A-D), although optical stimulation in probes with 60 μm windows often resulted in more intense spiking activity, presumably due to larger volume of activation (Figure 4).

**Figure 6:**
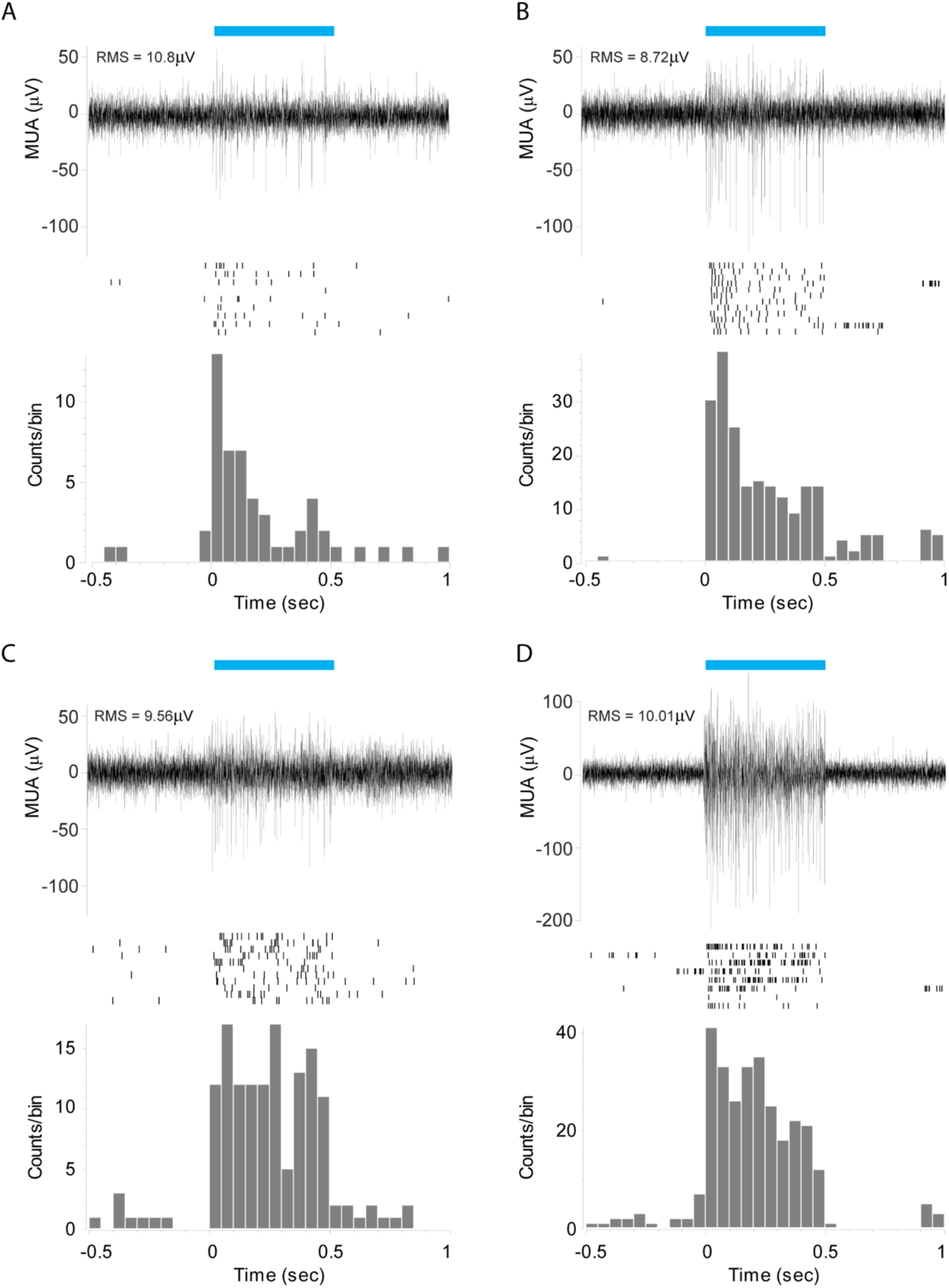
*In vivo* electrical recordings, striatum. (A) Top - Representative trace and raster plot of action potentials recorded in striatum of Adora2A-Cre x ChR2f/f mouse n=3, with a fibertrode containing 7μm electrode pad and 20μm optical window in response to continuous 500ms optical pulse. Bottom – Average peristimulus histogram of the number of action potentials. (B) C) and (D) same as (A) for a 7.5μm electrode pad and 40μm window, 7.5 μm electrode pad and 60μm window and 30μm with a 60μm window respectively. Plots represent recordings in 1 location 9-14 sweeps each, histogram bin = 50ms, RMS was calculated by averaging baseline of all traces.

To address whether fibertrodes can achieve short latency, pulse-locked optogenetic stimulation was performed with simultaneous recording of neural activity in layer 5 (L5) somatosensory cortex of Thy1-ChR2 mice. Compared to striatal SPNs, L5 pyramidal cells have faster spiking kinetics that enables better characterization of light-evoked action potentials. Short (2-5ms) optical pulses resulted in reliable stimulation of Thy1-ChR2 neurons (Figure 7A-B).

**Figure 7:**
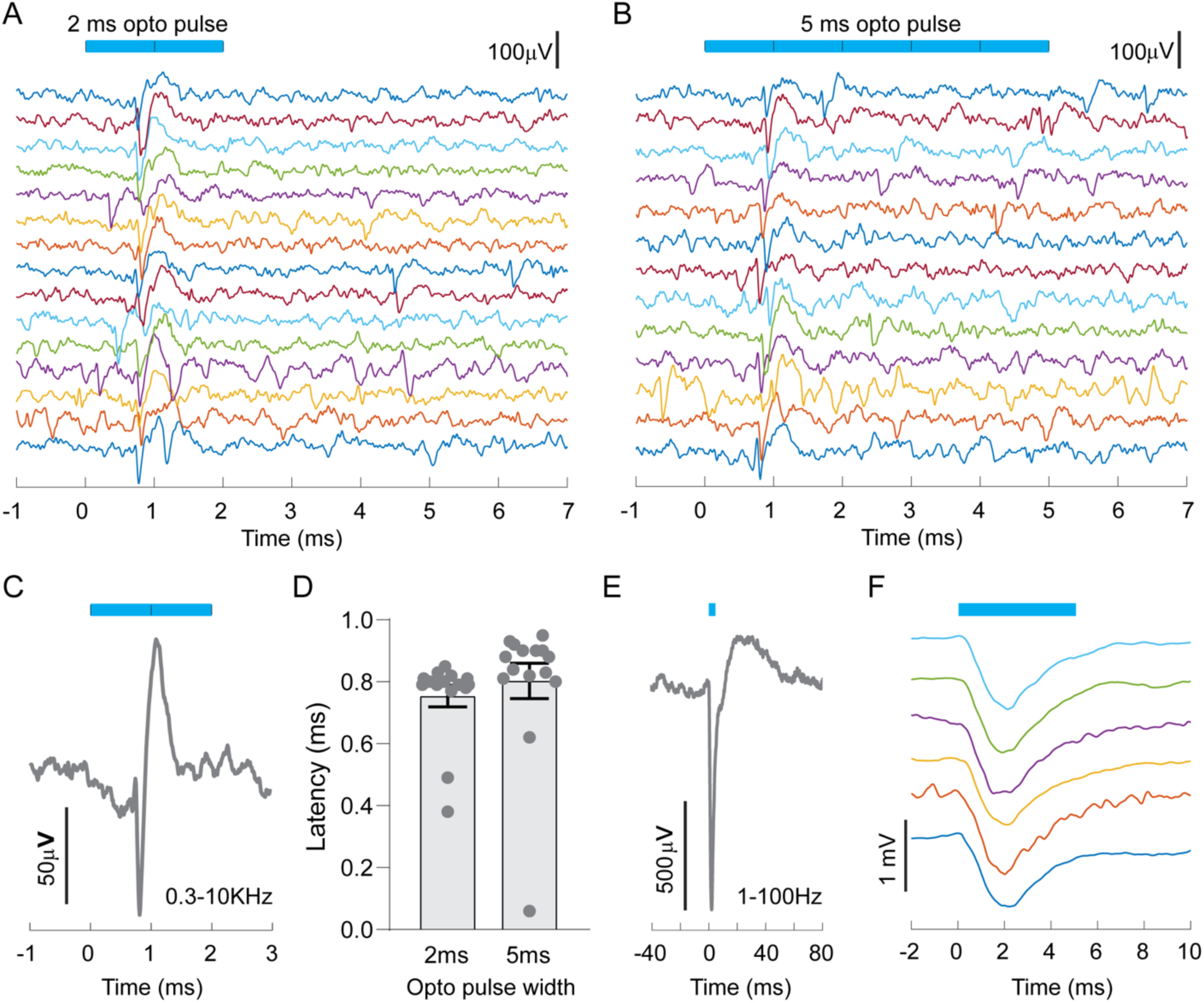
*In vivo* electrical recordings, cortex. (A) Electrophysiological traces of HPF neural activity recorded in somatosensory cortex of Thy1-ChR2 mouse in response to 2ms optical pulse. Note the reliable stimulation of action potentials in recorded neuron. (B) Optogenetic stimulation of action potentials in the same neuron recorded in (A) in response to 5ms optical pulse. Note the similar firing latency and absence of photoelectric artifacts at the onset and termination of optical pulses in both conditions. (C) Average action potential waveform of recordings depicted in A) aligned to optogenetic stimulus onset. Average of of 15 sweeps recorded with 2ms stimulation. (D) Variable latency of action potential stimulation induced by 2ms and 5ms optical pulse for the same recorded neuron indicate that waveform is not due to optical photoelectric artifacts. Slightly longer latency in response to 5ms stimuli is likely due to increased opsin desensitization compared to 2ms pulses. Analysis (mean +/-SEM) performed in Graph Pad Prism. (E) Local field potential deflection recorded in response to 5ms optical stimulation, average of 6 traces shown in F. (F) Individual traces of average LFP response depicted in E.

We did not detect photoelectric artefacts at the onset and offset of light stimulation (Figure 7A-B). Moreover, putative spikes occurred approximately 0.8ms after light onset (Figure 7C) and exhibited variable latency between stimulations (Figure 7D), indicating that they correspond to action potentials of stimulated cells rather than light-locked artefacts. LFPs recorded in response to 473nm light pulses (duration 5ms, power density ~10mW/mm^2^) delivered through the optical window elicited large LFP defections, indicating that fibertrodes have the capability to recruit large neuronal ensembles when used in higher light-power regimes. Despite this strong activation we did not detect optical artefacts in the LFP signal (Figure 7E-F), in agreement with *in vitro* experiments, where no light-induced electrical artefacts are detected for short light stimuli in the LF channel.

## DISCUSSION

Here we show that the photonic properties of tapered “fibertrodes” enable artefact-free extracellular recordings of neural activity during optogenetic stimulation of spatially confined neuronal populations. The use of a metal coated tapered fiber allowed the integration of an extracellular Pt electrode just above a light-emitting optical window. The effect of the taper generates an angled emission from the aperture that minimizes light incidence on the electrode contact, largely eliminating photoelectric artefacts arising from direct electrode illumination. While *in vitro* characterization of fibertrode electrical behavior during long light stimuli shows a small residual light-triggered artifact in the LFP frequency band (time constant τ>90ms), this is not detectable during LFP *in vivo* recordings, where optogenetic stimulation is triggered by brief light pulse-widths.

Our device allowed reliable optogenetic activation of ChR2-expressing neurons in both the striatum and cortex with high temporal precision. Importantly, because the optical window is placed in close proximity to the electrode contact, it is possible to achieve optogenetic activation at very low power regimes. By tailoring the window size and electrode size, fibertrodes can provide stimulation of tissue volumes ranging from 10^−^ 5 mm^3^ to 10^−3^ mm^3^ (Fig.4D), corresponding to ~1-100 neurons assuming a density of 10^5^ neurons/mm^3^. Fibertrodes thus offer the unprecedented possibility of spatially confining optogenetic stimulation to very restricted neuronal populations, while recording neuronal responses from or near that same volume. Other technologies are limited in this respect. For example, μLEDs are constrained by Lambertian emission profile and by electrical coupling between driving and readout channels[34], while flat-cleaved optical fibers have very low power density (2mW/mm^2^) that necessarily generates a larger excited volume, ~7×10^−3^ mm^3^ [21]. In addition, the highly localized, nanometric resolution of FIB milling and IBID processing on a stack of aluminum and Prl-C layers allows for the fabrication of probes with customized placement of electrode-window pairs anywhere along the taper. This level of high-resolution conformal patterning on non-planar surfaces has not been demonstrated yet for transparent conductive materials like ITO, PEDOT or graphene, that have been suggested to reduce photoelectric artefacts [9]. The size and shape of the fibertrode optical window can also be modified to illuminate specific volumes of tissues, and the impedance of the Pt electrode can be controlled by changing its surface area, even though it is realized on a highly non-planar fiber surface.

Moreover, the design concepts introduced in this work can be extended to other fibertrode configurations, for example by designing probes for wide-volume light delivery. In this case, widely-extended illumination could potentially impinge on the electrode, causing an electric artefact. A configuration that solves this problem is schematized in Supplementary Figure1A, whose light emission patterns are shown in Supplementary Figure 1B. Here, the recording site is placed just above the first emission diameter of the TF, and shielded from internal irradiation by the fiber cladding, which is preserved along the taper of 0.66NA borosilicate-glass fibers [22].

Representative *in vitro* (PBS) recordings in both HF and LF band show no photoelectric artefacts for 0.66NA even for 100ms light pulses (SupplementaryFigure1C, right). With this dielectric shielding, indeed, surface current density induced by propagating and evanescent modes does not arise, because the confinement is purely dielectric. The shielding effect of the cladding is supported by experiments with 0.39NA tapered fibers which instead lose their cladding during the heat and pull process and show conspicuous photoelectric artefacts (Supplementary Figure 1C, left).

In conclusion, the fibertrode presented in this work can enable new experiments for closed-loop optogenetic investigation of brain circuits on small and customizable brain volumes. More complex optrode system could be designed by arranging multiple fibers in customized arrays by developing fabrication strategies to pattern the fiber surface with multiple electrodes and connection lines; with on-demand light-emission patterns.

Lastly, TFs were recently employed for fluorescence light *collection in vivo* [35] potentially allowing fibertrodes to record both extracellular electric signal and fluorescence signals from genetically encoded functional indicators, such as GCaMP and dLight [26], [36]–[39]. Dual-mode recordings of this sort would greatly expand the range of fibertrode application, such as would allow correlations to be made between electrophysiology signals and functional fluorescence related to cellular events or neurotransmitters’ concentration. In conclusions fibertrodes, and the technologies behind their fabrication, will encompass the panel of functions required to close the loop in optogenetics allowing for optical activation, electrical readout and fiber photometry of functional neural activity.

## MATERIALS AND METHODS

### Animals

All experimental manipulations on mice were performed in accordance with protocols approved by the Harvard Standing Committee on Animal Care and guidelines described in the US National Institutes of Health *Guide for the Care and Use of Laboratory Animals*. For electrophysiological recordings in striatum *Adora2a*-Cre transgenic mice (GENSAT #KG139Gsat) were bred to conditional channelrhodopsin-2 (ChR2) expressing mice expressing ChR2(H134R)-EYFP under control of an upstream *loxP*-flanked STOP cassette (Ai32; referred to as *ChR2*^*f/f*^; The Jackson Laboratory #012569). Electrophysiological recordings in cortex were performed in Thy1-ChR2-YFP mice (The Jackson Laboratory #007612). In all experiments, male and female mice were used.

### Brain tissue processing and imaging

Recording locations were confirmed post hoc by whole brain sectioning and imaging. After recordings mice were deeply anesthetized with isoflurane and perfused transcardially with 4% paraformaldehyde in 0.1 M sodium phosphate buffer. Brains were fixed for 24 h at 4 °C, washed in phosphate buffer saline (PBS) and sectioned (100 μm) coronally using a vibratome (Leica VT1000s). Brain sections were mounted on glass slides, dried and mounted with ProLong antifade reagent containing DAPI (Molecular Probes). Whole brain sections were imaged with an Olympus VS110 slide-scanning microscope.

### Surgeries and in vivo recordings

For head bar implants and craniotomies mice were anesthetized with isoflurane and placed in a stereotaxic apparatus. After surgical removal of scalp and cleaning of the skull with saline and 70% ethanol, two craniotomies were made with a 0.5-mm burr micro drill at (AP-0 mm; ML-2.0 mm and AP-0.3 mm; ML-+1.5 mm; from Bregma) and sealed with Kwik-Cast silicone Elastomere. Animals were fitted with a custom-made titanium head bar using transparent glue (Loctite 454) and allowed to recover from anesthesia for 1 h on a heat pad at 38 °C. Following recovery from anesthesia, animals were head fixed and *in vivo* electrophysiological recordings were performed using different fibertrodes. Cortical recordings in L5 were done by positioning the electrode/window 1,250 μm deep from the brain surface, whereas striatal recordings were performed at a 2,500-2,750 μm depth. Optogenetic stimulation was achieved by coupling a 200μm core optical fiber (Thorlabs FT-200-UMT, 0.39 NA) using a 473-nm laser source. Light pulses were controlled with an Acousto-Optic Modulator (AA opto-Electronic) for fast shuttering and intensity control. Final light power was 2mW at the patchcord and pulse width was 2,5 or 500ms depending on experiments. All recordings were validated by post hoc serial histological analysis of electrode placement.

### Data acquisition and analysis

For electrophysiology recordings of *in vivo* neural activity extracellular signals were amplified and band-pass filtered (300Hz-10kHz) using a A-M Systems Model 1800 microelectrode AC amplifier (A-M Systems) and digitized at 40 kHz using a NI-DAQ 6363 acquisition board (National Instruments) and a custom version of ScanImage written in MATLAB (Mathworks). Off-line analysis of light evoked action potentials was performed using custom routines written in MATLAB. Spike sorting depicted in Figure 2 was performed via principal component analysis (PCA) in Offline Sorter v4 (Plexon). Waveform detection was performed after a fixed -50uV threshold detection followed and alignment by global minimum between sort start/stop times (1.5ms). PCA was performed on total pool of aligned waveforms and clustering identification performed manually. Spike waveform depicted in Figure 5 is the average trace of all evoked traces. Latency calculation was based on local minimum within 1ms of light onset. Variable latency results indicate that waveforms are not a result of photoelectric effect.

### Fibertrode fabrication process

The optrode is based on an optical fiber (0,39NA FT200UMT, Thorlabs) with one of the ends tapered down to a sub-micrometer tip with a low angle over a length of few millimeters. Tapered fibers where realized by the heat and pull method as described in[28]. An aluminum/Prl-C/aluminum stack (Fig. 1A) is deposited onto the taper by alternating thermal deposition and physical vapor deposition (STS Coating System) respectively. The fiber is kept under rotation during the first aluminum deposition to obtain metallization over the whole taper surface. The second aluminum layer is instead deposited without rotation of the fiber covering approximately half of the taper surface. Localized removal of the deposited materials to realize a squared optical aperture (Fig. 1B) is obtained by Focused Ion Beam milling (FEI® Helios™ NanoLab™ 600i DualBeam™, equipped with the Tomahawk FIB column). An insulated metallic wire with uninsulated ends is glued with a conductive epoxy to the metallized portion of the fiber a few millimiters apart from the tapered region. A second Prl-C layer is then deposited for final insulation of the probe. This process will also encapsulate the metal exposed at the optical window sidewalls during milling. Finally, the microelectrode is realized by FIB milling of the second Prl-C to expose the secondly deposited aluminum followed by Ion Beam Induced Deposition (IBID) of platinum to fabricate a circular microelectrode. Electrode is connected to the PCB by an insulated copper wire fixed at the external Al layer by means of Ag paste.

### Impedance characterization

Electrochemical Impedance Spectroscopy (EIS) was performed with a commercially available potentiostat system (ZIVE SP1, WonAtech Co., LTD.). Measurements were taken in phosphate buffered saline solution (PBS) in the three electrodes configuration, with the fabricated microelectrode, an AgCl wire and a Platinum wire immersed acting as the working electrode, the reference electrode and the counter electrode, respectively. Potentiostatic sinusoidal waveforms (10mV rms, 1-100 kHz) were applied with respect to the open-circuit potential (OCP) in order to let the actual surface conditions of the microelectrodes drive the measurement results.

### Fibertrode characterization

Light emission properties were assessed by injecting 473nm laser (Laser Quantum Ciel 473) into the optrode at different angles througt a galvanometric-mirror based scanning system as described in [22]. Optrodes were immersed in 30μM Fluorescein solution and light emission images were acquired using a fluorescence microscope (Scientifica Slicescope, 4X objective Olympus XLFLUOR4X/340 with immersion cap XL-CAP; sCMOS Hamamatsu ORCA-Flash4.0 V2 camera). Images were acquired for all fibertrodes. For each different input angle, optical output power was measured in air by placing the optical window in close proximity to a Thorlabs PM100USB power meter with S120VC sensor head. Power coupling efficiency was measured as the ratio between taper and patch fiber optical power output while power density was calculated as the ratio between the emitted light power and the window area (20μm^2^,40μm^2^ or 60μm^2^).

Devices were tested in PBS solution for light artefacts and electrical noise. 470nm light (Laser Quantum Ciel 473) was injected at full NA through a patchcord connected to the TF via a plastic sleeve. Electrode was connected to a 32-channel amplifier board (RHD2164 64-Channel Amplifier Board) and electrical recordings were performed using an Axon™pCLAMP™ Data Acquisition System and an Intan RHD2000 bradboard (20kHz sampling rate). pClamp software is used to trigger light stimulus delivered by 473 nm laser connected to one of the bradboard ADC channels, while the electrode is connected via a male connector to one of the headstage channels.

## ACKNOWLEDGEMENTS

B. Spagnolo, F. Pisanello F. Pisano acknowledge funding from the European Research Council under the European Union’s Horizon 2020 research and innovation program (#677683); M. Pisanello and M.D.V. acknowledge funding from the European Research Council under the European Union’s Horizon 2020 research and innovation program (#692943). L.S., M.D.V. and B.L.S. are funded by the US National Institutes of Health (U01NS094190). M. Pisanello, L.S., F. Pisanello, M.D.V., B.L.S. are funded by the US National Institutes of Health (1UF1NS108177-01). F. Pisanello and M.D.V. also acknowledge funding from the European Union’s Horizon 2020 research and innovation program under grant agreement (#828972). Authors also acknowledge Jaeeon Lee for help setting up the optrode fiber launch system.

